# Experimental evolution of symbiotic microbes without their partners can imply the presence of cooperative or antagonistic adaptations

**DOI:** 10.1101/2023.10.01.560375

**Authors:** Tyler J. Larsen, Cara Jefferson, Anthony Bartley, Joan E. Strassmann, David C. Queller

## Abstract

Microbes adapt to the presence of other species, but the fitness consequences of specific interactions are difficult to study in their natural context. We experimentally evolved symbiotic microbes in an artificial environment without access to the partners with whom they interact in nature. As organisms will tend to lose adaptations that they do not need due to drift or pleiotropic tradeoffs, we expect normally symbiotic microbes evolved in isolation to lose adaptations to help or harm their natural partners. The direction and magnitude of such changes can suggest whether the microbes had historically been selected to help or harm one another. We apply this method to the symbiosis between the social amoeba *Dictyostelium discoideum* and three intracellular bacterial endosymbionts, *Paraburkholderia agricolaris, P. hayleyella,* and *P. bonniea.* A minority of strains of *Paraburkholderia* and *D. discoideum* evolved differences in their effects on one another’s fitnesses, implying the existence of adaptations to one another that were lost when no longer relevant. Our results suggest that the degree to which *D. discoideum* and *Paraburkholderia* have adapted to help or harm one another can differ substantially between strains within each species, with some strains appearing to have a historically adversarial relationship, some strains a more benign relationship, and many strains no clear adaptations to one another at all. Our results underscore the complexity of microbial interactions in nature and suggest experimental evolution under relaxed selection is a potentially useful approach for studying adaptation in microbes.

## INTRODUCTION

Organisms cannot be understood without understanding how they interact with other species. Observing such interactions in microbes can be complicated because of the small scales involved (Young and Crawford 2004; Widder et al. 2016; Vos et al. 2013). Questions with answers that seem almost self-evident for larger organisms – such as ‘who interacts with whom?’, ‘is interacting costly or beneficial?’, and ‘are organisms adapted to help or harm other species?’ – can be very difficult to answer for microbes, particularly within their natural habitats. It is easy to see when a cheetah and a gazelle interact and what they are trying to do with respect to each other – they are enemies; their interactions are costly; they have adaptations to maximize their chances of success. Developing the same kind of understanding of the relationship between two soil microbes isolated from the same grain of soil – who might equally be inseparable friends, dire enemies, or neutral bystanders who never interact at all – can pose a special challenge.

One potentially complex relationship between microbes involves the social amoeba *Dictyostelium discoideum* and its bacterial symbionts *Paraburkholderia agricolaris, P. hayleyella,* and *P. bonniea* (Haselkorn et al. 2019). *D. discoideum* is a soil-dwelling amoeba that upon starvation aggregates into a multicellular fruiting body of durable spores that sit dormant and await dispersal atop a stalk of dead cells (Bonner 1944; Kessin 2001; Strassmann and Queller 2011). Three species of bacteria within the genus *Paraburkholderia* are capable of intracellularly infecting *D. discoideum* amoebae and then co-dispersing inside of and upon the surface of *D. discoideum* spores throughout multiple rounds of fruiting body formation (Haselkorn et al. 2019; DiSalvo et al. 2015; smith, Queller, and Strassmann 2014).

Laboratory experiments suggest that the interaction between *D. discoideum* and *Paraburkholderia* can have both positive and negative fitness consequences for each partner. *Paraburkholderia* infection usually reduces the fitness of *D. discoideum* hosts (DiSalvo et al. 2015), potentially because of the production of toxic compounds, the exploitation of intracellular host resources, or the disruption of normal host digestion. Despite this, *Paraburkholderia* infection also imbues *D. discoideum* with the ability to carry a simple microbiome of other bacteria – including suitable prey bacteria – as passengers through its social stage (DiSalvo et al. 2015; Brock et al. 2011; Dinh et al. 2018), which can facilitate *D. discoideum’s* colonization of environments impoverished in food bacteria (Brock et al. 2011). *Paraburkholderia* infection may also bolster its host’s resistance to toxins (Brock et al. 2016) and serve to defend against exploitation by uninfected *D. discoideum* strains (Brock et al. 2013). In turn, *Paraburkholderia* living within a host presumably enjoys a resource-rich environment where the costs of competition are minimized, as well as a dispersal advantage from riding along within its much larger and more mobile host. Direct comparisons of *Paraburkholderia* fitness with and without *D. discoideum* in the laboratory yield mixed results (Garcia et al. 2019). While *Paraburkholderia* can survive and divide inside of *D. discoideum* cells, the presence of *D. discoideum* in liquid coculture has been shown to significantly suppress extracellular *Paraburkholderia* growth rate.

The context dependence of the harm or help that *D. discoideum* and *Paraburkholderia* experience due to their association make it unclear whether they should be considered friends, foes, both, or neither. Accordingly it is difficult to predict to what extent *D. discoideum* and *Paraburkholderia* may have evolved adaptations to help or harm one another. As with all microbes, it is challenging to directly observe *D. discoideum* and *Paraburkholderia’s* interactions in nature.

In this study, we attempt to characterize the relationship between *D. discoideum, P. agricolaris, P. hayleyella,* and *P. bonniea* using an experimental evolution approach. We took multiple strains of each species and experimentally evolved them in a laboratory environment without access to one another. Under these conditions, any selective pressures *D. discoideum* and *Paraburkholderia* ordinarily exert upon one another in nature should be relaxed, and any preexisting adaptations maintained by these pressures should tend to be lost.

When a preexisting selective pressure is removed, adaptations driven by selection on that pressure are likely to atrophy – organisms seem to ‘use it or lose it’ and become less well-adapted to environments in which they do not live (Lahti et al. 2009; Johnson, Lahti, and Blumstein 2012). This can be seen in vestigial or transient traits like the hindlimbs of cetaceans, reduced digits in birds and ungulate mammals, or the eyes of cave fish, all of which appear to serve little modern function but may be the remnants of adaptations to past environments where selective pressures were different (Darwin 1888, 2012; Sadier, Sears, and Womack 2022). Bacteria that live inside larger organisms, and in particular intracellular endosymbionts, experience high rates of decay and loss of genes that are essential for free-living bacteria but not necessary inside of a host (McCutcheon and Moran 2012; Smith et al. 2006). Trait losses have also frequently been observed in experimental evolution experiments, wherein animal or microbial populations are often passaged in much simpler environments than those in which they evolved (Lee and Marx 2012; Renda et al. 2015; Hoffmann et al. 2001; Holland and Rice 1999; Jaenike 1993; Velicer, Kroos, and Lenski 1998; Behe 2010; Nilsson et al. 2005).

By the same reasons, organisms adapt to selective pressures imposed by other organisms with which they interact and lose these adaptations when their biotic context changes (Badgett et al. 2002; Carthey and Banks 2014; Berger et al. 2007; Atkinson 2006). If evolution in isolation makes an organism more destructive to a partner upon reintroduction, it suggests that the ancestral organism began with adaptations that benefitted (or at least reduced harm to) its partner’s fitness that were lost when they were rendered irrelevant. Conversely, if evolved organisms become significantly less damaging to their partner’s fitness, it suggests that the ancestral organism began with adaptations that harmed their partner. The traits that an organism loses when evolved under relaxed selection can thus provide evidence for adaptation to a cooperative or antagonistic interaction. Helpful or harmful, an organism cannot lose adaptations it does not have.

By looking for evidence of adaptations lost under relaxed selection, we hope to gain useful insight about the balance of cooperation and antagonism between *D. discoideum* and *Paraburkholderia* in nature. We previously published a study that outlined these ideas and attempted to demonstrate them using *D. discoideum, P. agricolaris,* and *P. hayleyella* (Larsen et al. 2021). Our later sequence analysis revealed cross-contamination issues that called our specific results into question. This study is intended to replicate its predecessor while incorporating numerous improvements. We employed species-specific PCR screens and DNA microsatellite analyses throughout and after experimental evolution to verify that lines had not been contaminated. In addition, we expanded the original experiment with the inclusion of a third symbiont species (*P. bonniea*) as well as non-host strains of *D. discoideum*.

## METHODS

### Strain selection

We selected three pairs each of naturally-occurring *D. discoideum/P. agricolaris* partners (QS70/bQS70, QS159/bQS159, QS161/bQS161), *D. discoideum/P. hayleyella* partners (QS11/bQS11, QS21/bQS21, QS69/bQS69), and *D. discoideum/P. bonniea* partners (QS395/bQS395, QS481/bQS481, QS859/bQS859).

Additionally, we selected three *D. discoideum* isolates with no associated *Paraburkholderia* infection (QS6, QS9, QS18 – hereafter called ‘non-hosts’). As we have reason to believe our non-host strains are on average less intimate partners in nature than *D. discoideum/Paraburkholderia* symbiont pairs, we expected that inclusion of these strains would help validate our prediction that loss of symbiotic partners should drive trait loss.

In all cases, we cured *D. discoideum* of *Paraburkholderia* infection using antibiotics (see below) before use, then reinfected as needed for phenotypic assays to ensure a consistent infective dose.

### Culture conditions

We conducted all experiments using SM/5 media (Loomis and Sussman 1966) (2 g glucose (Fisher Scientific), 2 g BactoPeptone (Oxoid), 2 g yeast extract (Oxoid), 0.2 g MgCl_2_ (Fisher Scientific), 1.9 g KHPO_4_ (Sigma-Aldrich), 1 g K_2_HPO_5 (_Fisher Scientific), and for solid media 15 g agar (Fisher Scientific) per liter deionized water). To start a fresh culture of *D. discoideum*, we diluted spores from −80°C glycerol frozen stocks in KK2 buffer (2.25g KH_2_PO_4_ (Sigma-Aldrich) and 0.67g K_2_HPO_4_ (Fisher Scientific) per liter deionized water). We plated 1.0×10^5^ total spores onto an SM/5 plate along with 200µl *K. pneumoniae* food bacteria resuspended in KK2 to an OD_600_ of 1.5. To start a fresh culture of any bacterial strain, we streaked stocks from the −80°C freezer for isolation on SM/5 plates.

### Antibiotic curing of *D. discoideum*

In order to remove associated bacteria, 1.0×10^5^ *D. discoideum* spores were plated on SM/5 agar medium containing 30ug/mL tetracycline and 10ug/mL ciprofloxacin with 200µl of *K. pneumoniae* resuspended in KK2 to an OD_600_ of 1.5. We allowed plates to grow at room temperature under ambient light until formation of fruiting bodies (3-5 days). We collected spores as above, then diluted and plated again as above. We then collected spores and performed spot test assays (described in (Brock et al. 2011)) and PCR using *Paraburkholderia-*specific primers to verify successful curing.

### Experimental evolution

We separated natural pairs of *D. discoideum/P. agricolaris, D. discoideum/P. hayleyella,* and *D. discoideum/P. bonniea* from their partners as described above and experimentally evolved them isolated from their partners.

All lines were evolved in triplicate on SM/5 plates. Lines including *D. discoideum* were supplemented with 200µl of *K. pneumoniae* food bacteria resuspended in KK2 to an OD_600_ of 1.5 on SM/5 plates. We incubated all lines at room temperature under ambient light and transferred 0.5% of the population to fresh plates every 48 hours by first harvesting all spores and cells into 10mL KK2 buffer using gentle pipetting and scraping of the agar surface, thoroughly vortexing the resulting suspensions, diluting them 200-fold, and plating 100µl onto fresh plates. Plates containing *D. discoideum* were additionally supplemented with 200uL of *K. pneumoniae* suspension as a food source.

The 48-hour transfer interval was selected to preempt *D. discoideum’s* fruiting stage and avoid selection for non-fruiting cheaters observed in previous studies (Kuzdzal-Fick et al. 2011). Every fifth transfer, we froze 1mL of the undiluted suspension of harvested cells at −80°C with 60% glycerol to act as a frozen archive.

During experimental evolution, routine checks were done for contamination of bacterial lines. We extracted DNA from 100uL of the undiluted suspension of harvested cells by boiling and ran PCR using primers specific for *P. agricolaris* and *P. hayleyella* (Garcia et al. 2019). In a small number of cases, tested lines showed amplification of the incorrect species or did not show amplification of the correct species, suggesting contamination had occurred. Affected lines were discarded and resumed from the most recent frozen ancestors using the process described above. Following experimental evolution (after the 30^th^ transfer) *D. discoideum* lines were checked for cross-contamination using DNA microsatellite analysis. We extracted DNA from 100µl of the undiluted suspension of harvested cells using CHELEX resin beads and amplified using fluorescently tagged PCR primers specific to highly variable microsatellite loci known to differ in length between *D. discoideum* strains (Smith 2004). The resulting amplicons were analyzed to compare evolved strains to ancestors. In order to maximize the usefulness of both tests for contamination, transfers were completed in a predefined order such that no two lines of the same species were transferred sequentially, and replicate lines were transferred at different times with reagents replaced and the workstation cleaned in between each replicated set.

After all lines had evolved in isolation for 30 transfers (representing approximately 130 generations), we reintroduced evolved strains to their partners and used two assays to test how evolution in isolation had changed the fitness effects that strains imposed upon their partners and the susceptibility of strains to their partners’ reciprocal effects. In all assays, *D. discoideum* and *Paraburkholderia* were paired with their naturally occurring partners, which had originally been co-isolated from soil All *Dictyostelium* lines were cured before use (see above) and reinfected for each assay to ensure consistency. For strains that did not have naturally occurring partners (the three non-host *D. discoideum* strains), we used the *Paraburkholderia agricolaris* strain bQS70 as a control partner in our experiments.

### Assays of *D. discoideum* spore production

To observe how experimental evolution of bacteria affected their impact on *D. discoideum* fitness, we compared total spore production of ancestral *D. discoideum* when uninfected, when infected by ancestral bacteria, and when infected by bacteria evolved without access to their hosts.

In order to observe how experimental evolution of *D. discoideum* affected its susceptibility to the effects of bacterial infection, we compared total spore production of ancestral *D. discoideum* and *D. discoideum* evolved in isolation when uninfected and when infected by ancestral bacteria.

For each spore production assay, we first suspended spores of each *D. discoideum* line of interest in KK2 buffer to a concentration of 10^6^ spores/mL. We suspended *K. pneumoniae* food bacteria and each bacterial symbiont line of interest in KK2 buffer at an OD_600_ of 1.5. We inoculated SM/5 plates with 100µl of the appropriate *D. discoideum* suspension (∼10^5^ spores) and 200µl of either 100% *K. pneumoniae* suspension or a 95:5 (vol:vol) mixture of *K. pneumoniae* and the appropriate bacterial symbiont suspension. We incubated plates at room temperature under ambient light for 5 days, after which we harvested all spores by washing plates with 10mL of a detergent solution of 0.1% NP40 in KK2. We diluted the resulting spore suspensions and counted on a hemocytometer to estimate total spore production across the entire plate. We performed each assay three times.

### Assays of bacterial growth rate

In order to observe how experimental evolution of *D. discoideum* affected its impact on bacterial symbiont fitness, we compared growth rates of ancestral bacteria when cultured alone and when co-cultured with ancestral *D. discoideum* or *D. discoideum* evolved in isolation.

In order to observe how experimental evolution of bacterial symbionts affected their susceptibility to the presence of *D. discoideum,* we compared growth rates of ancestral bacteria and bacteria evolved in isolation when cultured alone and when cocultured with ancestral *D. discoideum*.

For each growth rate assay, we inoculated each bacterial line of interest from freezer stocks into 2mL liquid SM/5 media and grew them to stationary phase overnight at 30°C in a shaking incubator. We quantified bacterial suspensions using a spectrophotometer and diluted them to an OD_600_ of 0.1. We prepared *D. discoideum* amoeba suspensions by inoculating SM/5 plates with *D. discoideum* spores from freezer stocks and *K. pneumoniae* food bacteria, incubating for ∼48 hours at room temperature, and harvesting amoebae by washing plates with KK2. To remove residual *K. pneumoniae,* we washed the resulting *D. discoideum* suspensions 3 times by centrifuging at 300xG for 3 minutes then resuspending the pellet in KK2. After the third wash, we resuspended pellets in KK2 buffer containing 30ug/mL tetracycline and 10ug/mL ciprofloxacin for 1 hour. We then centrifuged and washed the antibiotic-treated suspensions 2 more times with fresh KK2, counted amoebae using a hemocytometer, and diluted them to 10^7^ amoebae/mL. We obtained growth curves from a Tecan Spark microplate reader. We prepared wells of a 96 well plate by combining 100µl SM/5 broth with either 10µl OD_600_ 0.1 *Paraburkholderia* suspension or 10µl KK2 and either 10µl 10^7^ amoebae/mL *D. discoideum* suspension or 10µl KK2. We took OD_600_ measurements every 15 minutes for 48 hours to produce a growth curve for each well. We fit growth curves and calculated maximum specific growth rates using the fitr script (https://github.com/dcangst) in R.

We used an identical process to measure how experimental evolution of bacteria affected their susceptibility to the effects of *D. discoideum* by comparing growth rates of ancestral and evolved bacteria in the presence and absence of ancestral *D. discoideum*.

### Statistical analyses

We analyzed results using random effect negative binomial models with replicate line as a random effect.

For data comparing the effects of ancestral and evolved bacteria on ancestral *D. discoideum* spore production, the final, AIC-minimized model was Sporeproduction ∼ Treatment*Strain + (1|Line).

For data comparing the susceptibility of ancestral and evolved bacterial growth rates to the presence of ancestral *D. discoideum,* the final, AIC-minimized model was Growthrate ∼ Treatment+Strain + (1|Line).

For data comparing the effects of ancestral and evolved *D. discoideum* on ancestral bacterial growth rates, the final, AIC-minimized model was Growthrate ∼ Strain + (1|Line).

For data comparing the susceptibility of ancestral and evolved *D. discoideum* spore production to the presence of ancestral bacteria, the final, AIC-minimized model was Sporeproduction ∼ Strain + (1|Line).

We performed analysis using R version 4.0.4 (Team 2013), the *lme4* package(Bates et al. 2015), the *MASS* package (Ripley et al. 2013), and the *emmeans* package(Lenth 2022).

## RESULTS

### Experimental evolution reduces some *Paraburkholderia* strains’ antagonistic effects on *D. discoideum* spore production

To measure how experimental evolution of *Paraburkholderia* affected its fitness effects on its *D. discoideum* host, we compared *D. discoideum* spore production when uninfected and when infected by ancestral *Paraburkholderia* or *Paraburkholderia* experimentally evolved in the absence of *D. discoideum* (Figure 2).

**Figure 1.**
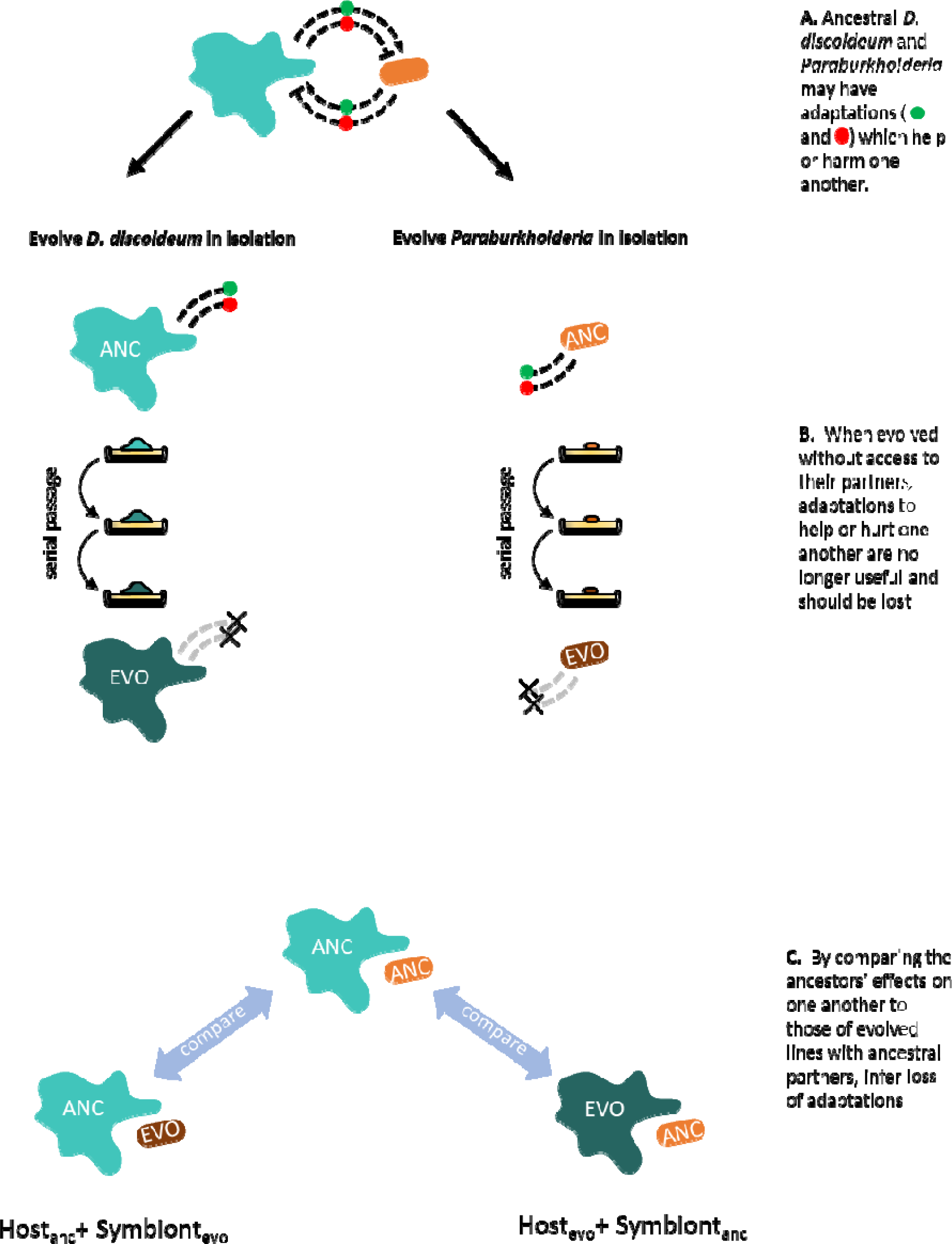
**using trait loss during experimental evolution to infer whether *D. discoideum* and *Paraburkholderia* have adaptations which help or harm one another in nature. A.** In nature*, D. discoideum* and *Paraburkholderia* potentially have adaptations which increase or decrease one another’s fitness. Here we represent these adaptations as secreted products (green or red circles) but any trait with an associated cost could be expected to follow the same logic. **B.** Regardless of whether they are helpful or harmful, these adaptations are costly or irrelevant in a laboratory environment where *D. discoideum* and *Paraburkholderia* do not have access to one another. **C.** Loss of beneficial or antagonistic adaptations can be inferred by reintroducing evolved *D. discoideum* or *Paraburkholderia* to their ancestral partners and looking for changes in the effects they have on their partners’ fitness.

**Figure 2.**
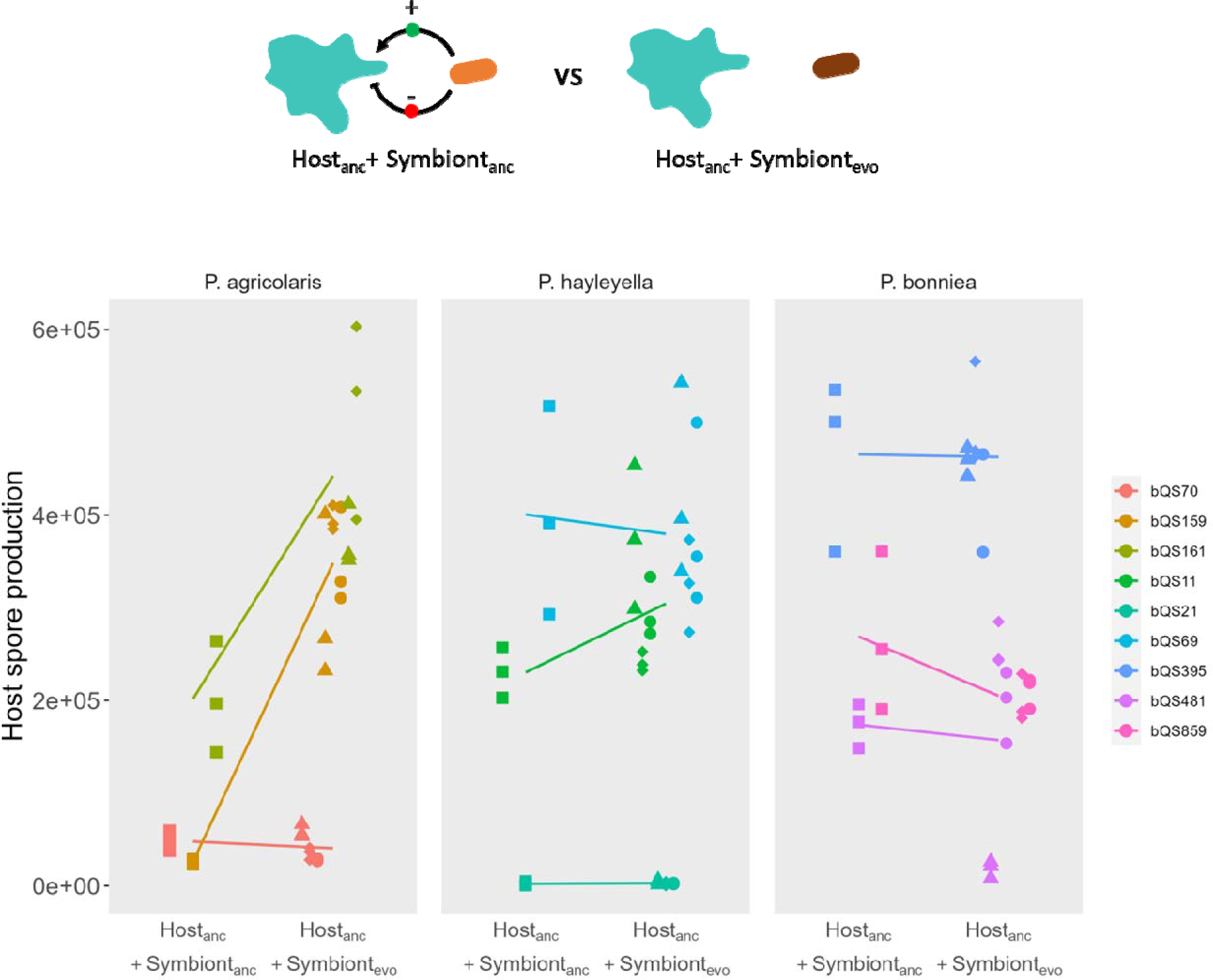
**Experimental evolution reduces some *Paraburkholderia* strains’ antagonistic effects on *D. discoideum* spore production** – Comparison of the effects of ancestral and evolved *Paraburkholderia* infections on ancestral *D. discoideum* spore production. Point shapes distinguish replicate evolved lines.

Overall, *Paraburkholderia* strains experimentally evolved in a host-free environment had a significantly reduced toxic effect on *D. discoideum* host spore production (β=-1.05, SE=0.126, p<0.0001, Table 1). When data for P. agricolaris, P. hayleyella, and P. bonniea were analyzed separately, only P. agricolaris showed significant reductions in toxicity. By contrast, the negative effects of strains of *P. hayleyella* and *P. bonniea* did not significantly change due to experimental evolution.

**Table 1.** Summary of statistical analysis.

|  |  |
| --- | --- |
| Analysis | **Figure 2** – Comparison of evolved and ancestral *Paraburkholderia*'s effects on host spore production |
| Model formula/R code | **Totalspores~Treatment\*Strain + (1\|Line)** |
| All data | Analysis of variance model comparison: Treatment $\chi^2_9=26.581$, $p=0.0016^*$. Strain $\chi^2_{16}=100.2$, $p<0.0001^*$<br>Emmeans effect of Treatment across all *Paraburkholderia* – $\beta=-1.05$, $SE=0.126$, $p<0.0001^*$ |
| *P. agricolaris* data only | Emmeans effect of Treatment across all *P. agricolaris* – $\beta=-1.05$, $SE=0.126$, $p<0.0001^*$<br>Emmeans bQS70 Ancestor vs Evolved – $\beta=0.219$, $SE=0.215$, $p=0.3090$<br>Emmeans bQS159 Ancestor vs Evolved – $\beta=-2.597$, $SE=0.215$, $p<0.001^*$<br>Emmeans bQS161 Ancestor vs Evolved – $\beta=-0.773$, $SE=0.228$, $p=0.0007^*$ |
| *P. hayleyella* data only | Emmeans effect of Treatment across all *P. hayleyella* – $\beta=-0.154$, $SE=0.121$, $p=0.2026$<br>Emmeans bQS11 Ancestor vs Evolved – $\beta=-0.263$, $SE=0.210$, $p=0.2105$<br>Emmeans bQS21 Ancestor vs Evolved – $\beta=-0.261$, $SE=0.210$, $p=0.2157$<br>Emmeans bQS69 Ancestor vs Evolved – $\beta=0.059$, $SE=0.210$, $p=0.7752$ |
| *P. bonniea* data only | Emmeans effect of Treatment across all *P. bonniea* – $\beta=0.285$, $SE=0.42$, $p=0.4968$<br>Emmeans bQS395 Ancestor vs Evolved – $\beta=0.008$, $SE=0.711$, $p=0.9910$<br>Emmeans bQS481 Ancestor vs Evolved – $\beta=0.575$, $SE=0.713$, $p=0.4196$<br>Emmeans bQS859 Ancestor vs Evolved – $\beta=0.272$, $SE=0.757$, $p=0.7191$ |
| Analysis | **Figure 3** – Comparison of evolved and ancestral *Paraburkholderia*'s susceptibility to host growth suppression |
| Model formula/R code | **Growthrate~Treatment\*Strain + (1\|Line)** |
| All data | Analysis of variance model comparison: Treatment $\chi^2_9=13.305$, $p=.1493$. Strain $\chi^2_{16}=42.797$, $p=0.0003^*$<br>Emmeans effect of Treatment across all *Paraburkholderia* – $\beta=-1.54$, $SE=0.784$, $p=0.0493^*$ |
| *P. agricolaris* data only | Emmeans effect of Treatment across all *P. agricolaris* – $\beta=-0.253$, $SE=0.112$, $p=0.0243^*$<br>Emmeans bQS70 Ancestor vs Evolved – $\beta=-0.152$, $SE=0.191$, $p=0.425$<br>Emmeans bQS159 Ancestor vs Evolved – $\beta=-0.447$, $SE=0.191$, $p=0.0191^*$<br>Emmeans bQS161 Ancestor vs Evolved – $\beta=-0.160$, $SE=0.202$, $p=0.428$ |
| *P. hayleyella* data only | Emmeans effect of Treatment across all *P. hayleyella* – $\beta=-1.28$, $SE=2.23$, $p=0.5666$<br>Emmeans bQS11 Ancestor vs Evolved – $\beta=-0.933$, $SE=3.43$, $p=0.7856$<br>Emmeans bQS21 Ancestor vs Evolved – $\beta=0.536$, $SE=3.86$, $p=0.8894$<br>Emmeans bQS69 Ancestor vs Evolved – $\beta=-3.431$, $SE=3.80$, $p=0.3660$ |
| *P. bonniea* data only | Emmeans effect of Treatment across all *P. bonniea* – $\beta=-2.99$, $SE=0.301$, $p<0.0001^*$<br>Emmeans bQS395 Ancestor vs Evolved – $\beta=-0.0269$, $SE=0.392$, $p=0.9453$<br>Emmeans bQS481 Ancestor vs Evolved – $\beta=-8.6176$, $SE=0.698$, $p<0.0001^*$<br>Emmeans bQS859 Ancestor vs Evolved – $\beta=-0.3197$, $SE=0.417$, $p=0.4428$ |
| Analysis | **Figure 4** – Comparison of evolved and ancestral *D. discoideum*'s effects on symbiont growth rate |
| Model formula/R code | **Growthrate~Treatment\*Strain + (1\|Line)** |
| All data | Analysis of variance model comparison: Treatment $\chi^2_{12}=23.044$, $p=0.0273^*$. Strain $\chi^2_{22}=64.279$, $p<0.0001^*$ |
| | Emmeans effect of Treatment across all *Dictyostelium* – $\beta=0.382$, SE=0.743, p=0.6076<br>Emmeans QS70 Ancestor vs Evolved – $\beta=-0.6024$, SE=2.25, p=0.7894<br>Emmeans QS159 Ancestor vs Evolved – $\beta=0.0228$, SE=2.52, p=0.9928<br>Emmeans QS161 Ancestor vs Evolved – $\beta=-0.0511$, SE=2.50, p=0.9837<br>Emmeans QS11 Ancestor vs Evolved – $\beta=-3.2982$, SE=2.53, p=0.1930<br>Emmeans QS21 Ancestor vs Evolved – $\beta=-0.7319$, SE=2.56, p=0.7754<br>Emmeans QS69 Ancestor vs Evolved – $\beta=-5.7621$, SE=2.62, p=0.0280*<br>Emmeans QS395 Ancestor vs Evolved – $\beta=11.198$, SE=2.63, p<.0001*<br>Emmeans QS481 Ancestor vs Evolved – $\beta=-5.7126$, SE=2.78, p=0.0402*<br>Emmeans QS859 Ancestor vs Evolved – $\beta=0.5607$, SE=2.52, p=0.8240<br>Emmeans QS6 Ancestor vs Evolved – $\beta=0.2180$, SE=2.57, p=0.9324<br>Emmeans QS9 Ancestor vs Evolved – $\beta=0.8118$, SE=2.56, p=0.7507<br>Emmeans QS18 Ancestor vs Evolved – $\beta=-0.1332$, SE=2.47, p=0.9570 |
| Analysis | **Figure 5** – Comparison of evolved and ancestral *D. discoideum*'s susceptibility to symbiont antagonism |
| Model formula/R code | **Totalspores~Treatment\*Strain + (1\|Line)** |
| All data | Analysis of variance model comparison: Treatment $\chi^2_{12}=9.4185$, p=0.6668. Strain $\chi^2_{22}=96.89$, p<0.0001*<br>Emmeans effect of Treatment across all *Dictyostelium* – $\beta=0.502$, SE=0.222, p=0.0241*<br>Emmeans QS70 Ancestor vs Evolved – $\beta=1.3476$, SE=0.774, p=0.0817<br>Emmeans QS159 Ancestor vs Evolved – $\beta=0.5616$, SE=0.737, p=0.4609<br>Emmeans QS161 Ancestor vs Evolved – $\beta=0.2673$, SE=0.765, p=0.7269<br>Emmeans QS11 Ancestor vs Evolved – $\beta=0.7442$, SE=0.765, p=0.3309<br>Emmeans QS21 Ancestor vs Evolved – $\beta=1.4355$, SE=0.773, p=0.0634<br>Emmeans QS69 Ancestor vs Evolved – $\beta=0.1793$, SE=0.767, p=0.8153<br>Emmeans QS395 Ancestor vs Evolved – $\beta=0.5036$, SE=0.770, p=0.5132<br>Emmeans QS481 Ancestor vs Evolved – $\beta=-0.1980$, SE=0.815, p=0.8081<br>Emmeans QS859 Ancestor vs Evolved – $\beta=-0.1534$, SE=0.762, p=0.8404<br>Emmeans QS6 Ancestor vs Evolved – $\beta=0.3464$, SE=0.756, p=0.6469<br>Emmeans QS9 Ancestor vs Evolved – $\beta=0.9021$, SE=0.763, p=0.2368<br>Emmeans QS18 Ancestor vs Evolved – $\beta=0.0838$, SE=0.770, p=0.9133 |

### Experimental evolution reduces some *Paraburkholderia* strains’ susceptibility to *D. discoideum* growth suppression

To measure how experimental evolution of *Paraburkholderia* affected its susceptibility to the growth suppression of co-cultured *D. discoideum,* we compared the growth rate of ancestral or evolved *Paraburkholderia* with and without the presence of ancestral *D. discoideum* (Figure 3).

**Figure 3.**
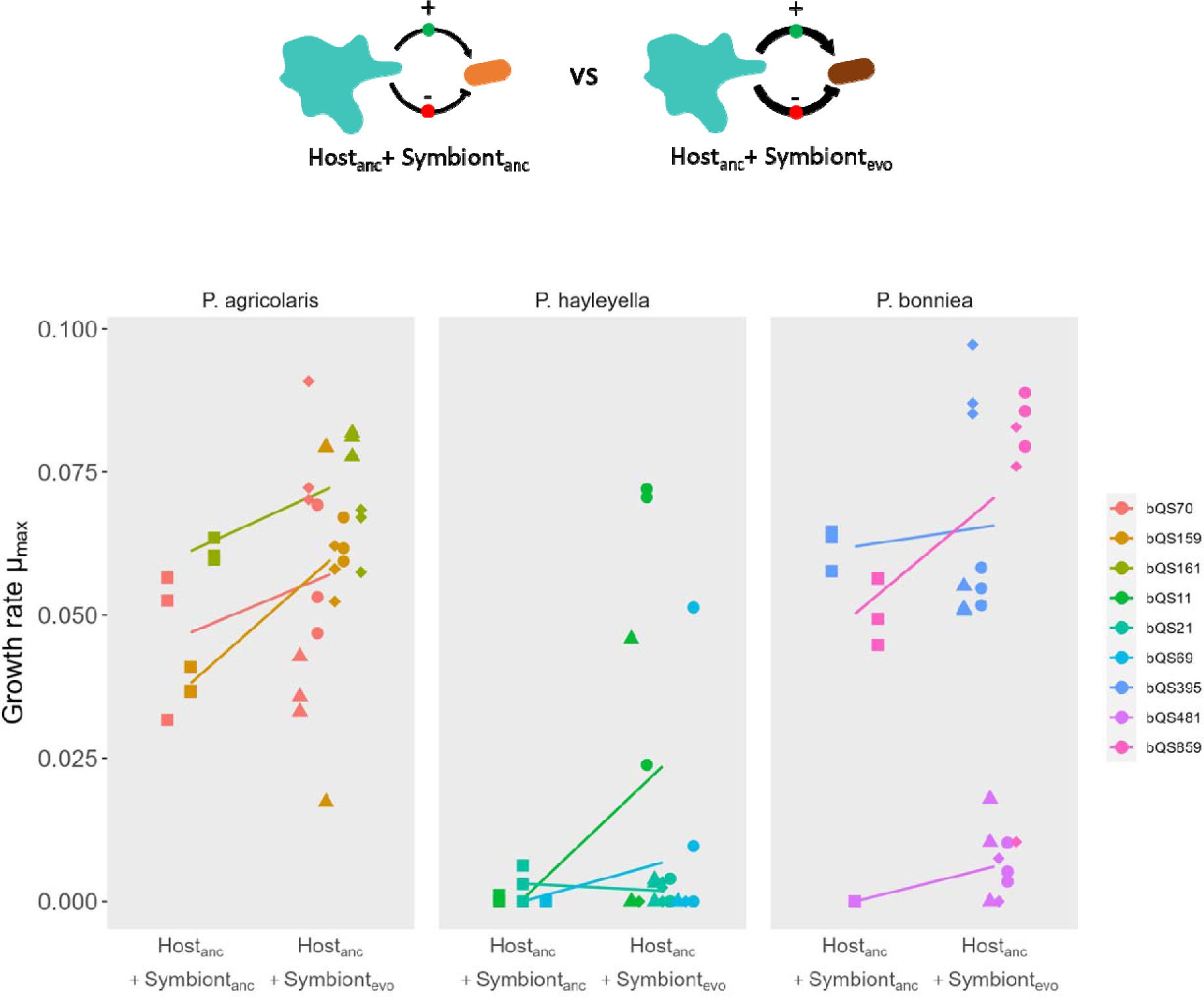
**Experimental evolution reduces some *Paraburkholderia* strains’ susceptibility to *D. discoideum* growth suppression –** Comparison of the effects of the presence of ancestral *D. discoideum* host on ancestral and evolved *Paraburkholderia* growth rates. Point shapes distinguish replicate evolved lines.

Overall, *Paraburkholderia* strains experimentally evolved in a host-free environment were significantly less susceptible to growth suppression by *D. discoideum* (β=-1.54, SE=0.784, p=0.0493, **Table 1**). When data for *P. agricolaris, P. hayleyella,* and *P. bonniea* were analyzed separately, *P. agricolaris* and *P. bonniea* had significantly increased susceptibility, though the effect in both species appeared to be the result of a single strain. *P. hayleyella’s* susceptibility did not significantly change due to experimental evolution.

### Experimental evolution did not significantly change *D. discoideum’s* antagonistic effects on *Paraburkholderia* growth rate

To measure how experimental evolution of *D. discoideum* affected its ability to suppress the growth rate of *Paraburkholderia* in its environment, we compared *Paraburkholderia* growth rates when grown alone and when co-cultured with either ancestral *D. discoideum* or *D. discoideum* experimentally evolved in the absence of bacterial symbionts (Figure 4).

**Figure 4.**
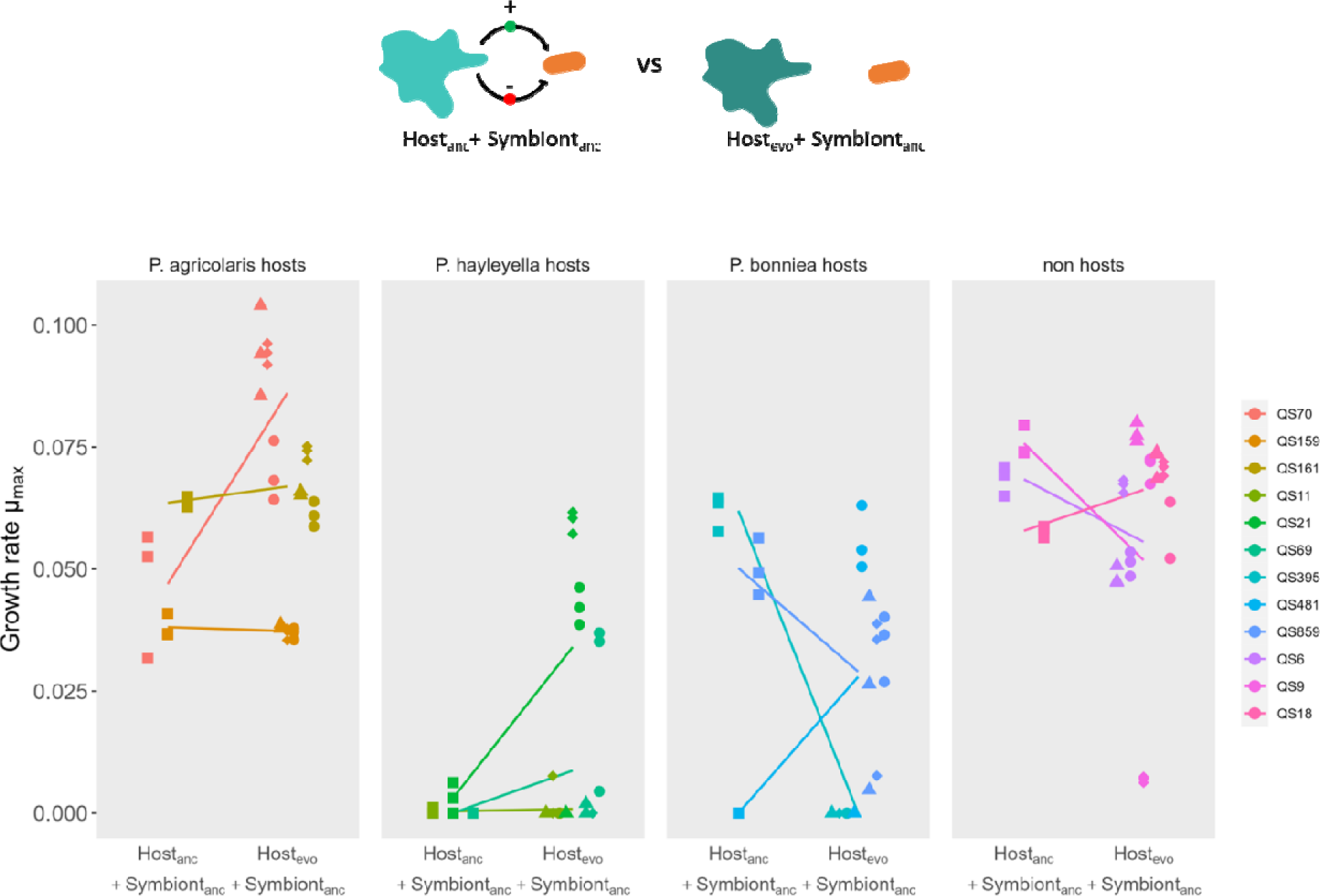
**Experimental evolution did not significantly change *D. discoideum’s* antagonistic effects on *Paraburkholderia* growth rate** – Comparison of the effects of the presence of ancestral and evolved *D. discoideum* on ancestral *Paraburkholderia* growth rates. Point shapes distinguish replicate evolved lines.

We did not observe a statistically significant overall effect on *D. discoideum’s* effect on *Paraburkholderia* growth rate (β=0.382, SE=0.743, p=0.6076, **Table 1).** When strains were considered individually, two strains were found to have evolved significantly reduced growth suppression and one strain was found to have evolved significantly increased growth suppression, relative to their respective ancestors.

### Experimental evolution increases some *D. discoideum* strains’ susceptibility to *Paraburkholderia* toxicity

To measure how experimental evolution of *D. discoideum* affected its susceptibility to the negative fitness consequences of *Paraburkholderia* infection, we compared the spore production of infected and uninfected ancestral *D. discoideum* or *D. discoideum* evolved in the absence of bacterial symbionts (Figure 5).

**Figure 5.**
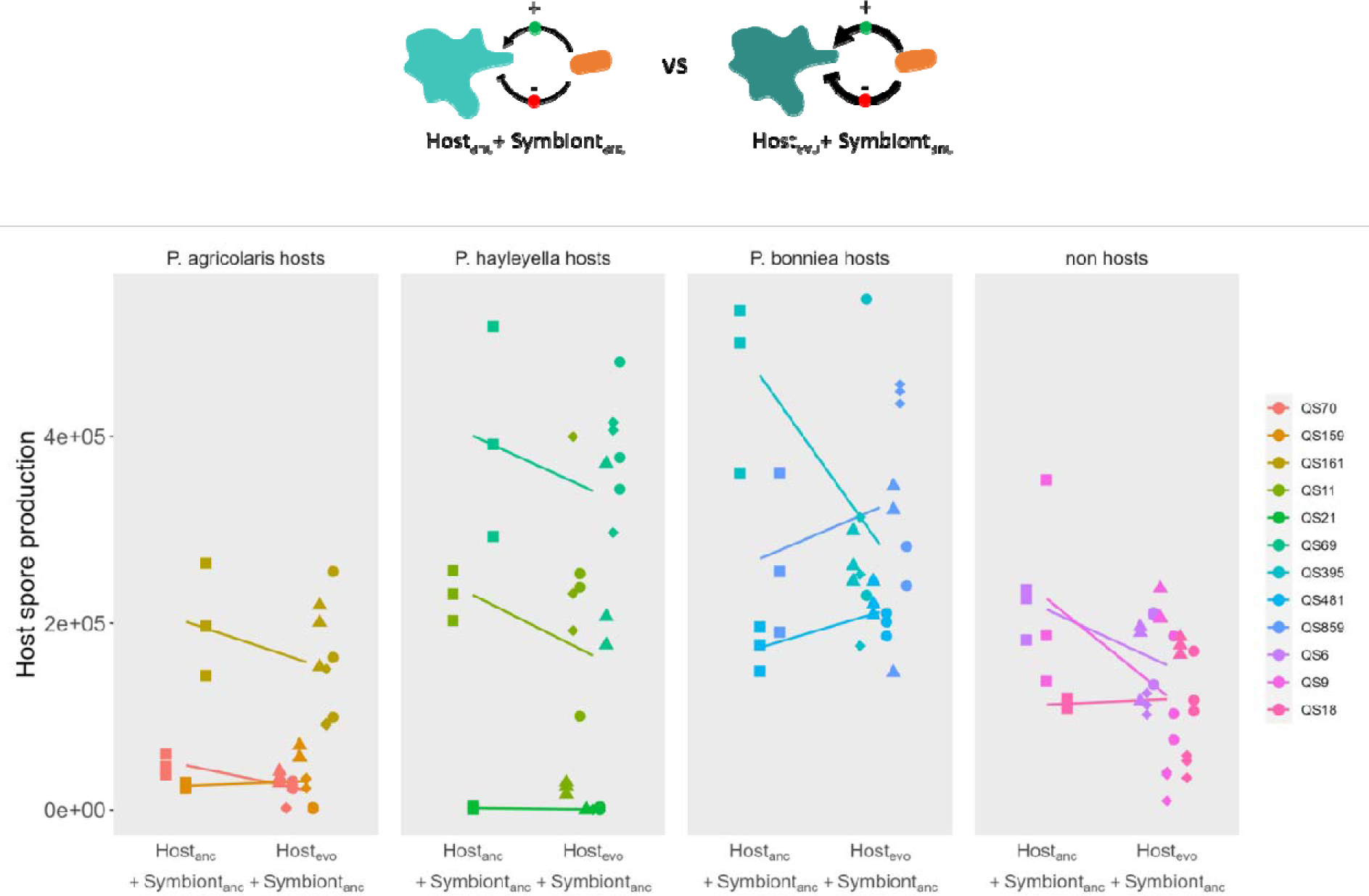
**Experimental evolution increases some *D. discoideum* strains’ susceptibility to *Paraburkholderia* toxicity** – Comparison of the effects of ancestral *Paraburkholderia* infections on ancestral and evolved *D. discoideum* spore production. Point shapes distinguish replicate evolved lines.

Overall, *D. discoideum* strains experimentally evolved in a symbiont-free environment were significantly more susceptible to the toxic effects of ancestral *Paraburkholderia* infection (β=0.502, SE=0.222, p=0.0241, **Table 1)**. We did not observe significant effects of experimental evolution for any strain considered individually.

## DISCUSSION

In a laboratory environment where they have no access to one another, *D. discoideum* and *Paraburkholderia* should lose adaptations they have to help or harm one another in nature. In this study, we sought to use this prediction to infer the selective pressures *D. discoideum* and *Paraburkholderia* impose upon one another in nature. Rather than ask how microbes adapt to novel artificial selective pressures as in many experimental evolution studies, we studied how *D. discoideum* and *Paraburkholderia* lose adaptations when selective pressures they experience in nature are taken away.

We found a statistically significant overall effect of experimental evolution on *Paraburkholderia’s* toxic effects on *D. discoideum* hosts’ spore production, such that ancestral *D. discoideum* infected by evolved *Paraburkholderia* produced more spores than ancestral *D. discoideum* infected by ancestral *Paraburkholderia* (Figure 2). This reduction in *Paraburkholderia* toxicity can be interpreted as evidence that the wild *Paraburkholderia* strains with which we started the experiment had adaptations that made them toxic in the first place. When moved to the laboratory where selective pressures are relaxed, these adaptations may have become neutral or even maladaptive and thus evolved away over time. Thus, the results of this assay imply an antagonistic relationship between these strains and their hosts. However, the magnitude and direction of the effect of experimental evolution varied considerably between strains, with most of the statistically significant overall effect being driven by the *P. agricolaris* strains bQS159 and bQS161. Other strains – even other strains within the same species, like bQS70 – showed no change or even an increase in toxicity, implying that the nature of the relationship between *D. discoideum* and *Paraburkholderia* may depend on the specific strain combinations involved.

We detected a small but significant overall effect on the susceptibility of experimentally evolved *Paraburkholderia* to growth suppression by *D. discoideum,* such that experimentally evolved *Paraburkholderia* grew faster in the presence of ancestral *D. discoideum* than did ancestral *Paraburkholderia* (Figure 3). Conversely, two strains – *P. agricolaris* strain bQS70 and *P. bonniea* strain bQS481 – evolved to become significantly *less* susceptible to growth suppression when no longer exposed to their hosts. This somewhat counterintuitive result may reflect the loss of adaptations with which they might normally capitalize on *D. discoideum’s* presence. Most *Paraburkholderia* strains, however – in particular strains of *P. hayleyella* and *P. bonniea* – did not evolve significantly different toxicity or susceptibility when evolved without access to their hosts. This negative result may indicate a lack of adaptations specifically evolved to help or harm *D. discoideum* in most strains. Notably, *P. hayleyella* and *P. bonniea* have substantially reduced genomes compared to other *Paraburkholderia* species (Brock et al. 2020). While small genomes are often associated with an obligately endosymbiotic lifestyle (McCutcheon and Moran 2012), they may also reduce the evolvability of microbes – it may be that *P. hayleyella* and *P. bonniea* do have adaptations to help or harm *D. discoideum,* but that they are more evolutionarily constrained and so did not lose these traits during the duration of our experiment.

Our results for *D. discoideum* also indicated important variation between strains. Two strains of *D. discoideum* – QS70 and QS481 – evolved significantly reduced toxic effects on the growth rate of *Paraburkholderia* symbionts in co-culture (Figure 4). As with the *P. agricolaris* results, these results imply that these *D. discoideum* strains possessed adaptations to harm *Paraburkholderia* that were lost when there were no longer *Paraburkholderia* present to harm. By contrast, QS395 evolved to become significantly *more* toxic, implying the loss of adaptations to increase *Paraburkholderia* growth rate. As with the *Paraburkholderia* results, however, most *D. discoideum* strains did not evolve significantly different toxicity nor susceptibility to *Paraburkholderia* toxicity (Figure 5), implying that these strains may lack adaptations to help or harm *Paraburkholderia* in nature. *D. discoideum* has been observed to have an uncommonly low mutation rate (Saxer et al. 2012), which may explain our generally negative results, though past experimental evolution experiments in *D. discoideum* (Kuzdzal-Fick et al. 2011; Larsen et al. 2021) lead us to believe the timeframe of this study was long enough to result in observable changes. Alternatively, the lack of evidence for adaptations affecting *Paraburkholderia* fitness in most *D. discoideum* strains in our study may reflect a lack of adaptations *specific* to *Paraburkholderia.* Under the conditions of our experimental evolution, *D. discoideum* did not have access to *Paraburkholderia,* but did have access to the prey bacterium *Klebsiella pneumoniae.* To the extent that *D. discoideum* might influence symbiont and prey fitness via the same mechanisms, the presence of food bacteria may have resulted in the maintenance of selective pressure for these mechanisms to remain throughout our experiment.

Our results suggest that whether the relationship between *D. discoideum* and its *Paraburkholderia* symbionts is cooperative, antagonistic, or neutral in nature may vary considerably even among closely related strains. We observed relatively few statistically significant overall or species-level trends. Instead, different strains of the same species sometimes showed markedly different degrees of initial antagonism (for example, see Figure 3.2C) and sometimes opposite responses to experimental evolution. Though the strains we employed in this study were isolated at roughly the same site at roughly the same date, the apparent differences in the degree to which they have adapted to cooperate or antagonize their partners may reflect adaptation to conditions varying over smaller scales of space and time. The relationship between *D. discoideum* and *Paraburkholderia* is sensitive to environmental conditions, and if different strains experienced slightly different conditions, they might well adapt in different directions or to different degrees.

Even traits with clear adaptive function come with associated costs, whether in the form of energetic costs of the traits themselves or more complicated tradeoffs among traits (Guillaume and Otto 2012; Rodríguez-Verdugo et al. 2014). A microbe living in nature must make compromises between mutually incompatible traits impacted by multiple selective pressures. When the microbe is moved to a simpler laboratory environment with fewer selective pressures, some compromises will no longer be necessary and traits should be lost to reflect that. In principle, it is possible that pleiotropic effects could instead result in gains of functionality (Meyer et al. 2010), but these situations should be the exception. Most pleiotropy will be negative rather than positive for the same reason that most mutational effects are negative – there are more ways to break complex adaptations than there are to enhance them (Johnson, Lahti, and Blumstein 2012). Given time, geographically isolated populations tend to lose reproductive compatibility – effectively a loss of function – rather than increase it, and symbionts living in a simplified host environment tend to lose adaptations to their prior environments rather than enhancing them(Dodd 1989; McCutcheon and Moran 2012; Campbell et al. 2015).

Our results provide new insights into *D. discoideum* and *Paraburkholderia’s* relationship in nature. Most of all, they suggest that no one-size-fits-all description can be made, and that just as the relationship between *D. discoideum* and *Paraburkholderia* can be very different under different environmental contexts, it can also be very different depending on the specific strain(s) being considered. The different outcomes we observed even between closely related strains emphasize the need to account for variation within species when studying wild microbes and the risk of overattributing results derived from observations of only one or a few strains. The approach we have taken in this study takes advantage of the differences between natural and laboratory environments by looking for traits lost when selective pressures are relaxed. Similar approaches could usefully supplement other studies of adaptation, particularly those performed using microbes that do not have long histories of use in the laboratory and for which the natural context may be especially unclear or hard to recreate.

